# Non-vocal motor deficits in a transgenic mouse model linked to stuttering disorders

**DOI:** 10.1101/2025.08.08.669441

**Authors:** Marissa Millwater, Maximillian Weinhold, Camryn Bragg, Harjot Kaur, Ruli Zhang, Shahriar SheikhBahaei

**Affiliations:** Neuron-Glia Signaling and Circuits Unit, National Institute of Neurological Disorders and Stroke (NINDS), National Institutes of Health (NIH), Bethesda, 20892 MD, USA; Center for Nervous Systems Disorders, Department of Neurobiology and Behavior, Stony Brook University, Stony Brook, 11794 NY, USA

**Keywords:** astrocytes, breathing, brainstem, grooming, cortico-striatal circuits

## Abstract

Stuttering is a neurodevelopmental disorder characterized by involuntary disruptions in speech. In addition, non-vocal motor impairments are reported in some individuals who stutter. Although its precise cause remains unknown, mutations in lysosomal trafficking proteins (such as GNPTAB) have been identified in a subgroup of people who stutter. To understand the functional significance of these mutations, transgenic *Gnptab* mice have been developed, and as expected, these mice exhibit vocal deficits throughout developmental stages. However, whether these mice also display non-vocal motor impairments is unknown. Our data reveal deficits in the breathing, locomotion, and grooming behaviors of the *Gnptab* mouse model, outlining a broader phenotype linked to *GNPTAB* mutations in stuttering. These findings suggest that lysosomal dysfunction may disrupt astrocyte-regulated motor circuits, affecting both vocal and non-vocal rhythmic behaviors that are central to stuttering neurophysiological symptoms.

## INTRODUCTION

Developmental stuttering, or childhood-onset fluency disorder, persists into adulthood for many and is estimated to affect over 80 million adults worldwide [1], [2]. Although the underlying mechanism(s) are not fully understood, stuttering is known to be a neurodevelopmental disorder to which genetic substrates contribute significantly [3], [4].

Existing data from people who stutter shows altered connectivity in brain regions controlling speech [5] or reduced beta band power in resting-state electroencephalography [EEG; [6]]. Though many investigations remain underway, at large, these investigations suggest a broader change in brain circuits that could involve a spectrum of motor dysfunctions extending beyond stuttering phenotypes. Non-vocal motor symptoms, including hyperactivity, differences in sequential finger movements, and breathing phenotypes, have been reported in some people who stutter [7]–[11] [12][13], [14]. However, the circuits and cellular mechanisms underlying these symptoms are not yet fully understood. Animal models have proven invaluable in understanding the neurogenetic mechanisms underlying motor behaviors at the circuit and cellular levels [15], including those involved in stuttering disorders [2], [16]–[19].

Here, in a transgenic mouse with a mutation in the *Gnptab* gene (*Gnptab*-mutant), which is involved in lysosomal targeting pathways and is associated with stuttering in humans, we investigated non-vocal motor behaviors (i.e., breathing and self-grooming). Our data indicate that *Gnptab*-mutant mice exhibit deficits in these non-vocal motor behaviors in addition to the previously reported vocal phenotypes [20]. When the mutation was engineered exclusively in astrocytes, mice displayed similar breathing and grooming phenotypes, suggesting that astrocytes may play a significant role in the development of non-vocal motor symptoms in stuttering. Our data adds to the accumulating evidence [17]–[19], [21] suggesting that *Gnptab*-mutant mice provide a framework to study the developmental mechanisms underlying motor speech disorders at the circuit and cellular levels.

## METHODS

### Animals

Adult male mice (3-5 months old) were used in this study. All procedures were performed in accordance with the *Guide for the Care and Use of Laboratory Animals* and were approved by the institutional Animal Care Use Committee. Mice were housed in a temperature and humidity-controlled facility with a normal light-dark cycle (12h:12h). Access to food and water was provided *ad libitum*.

### Measurement and analysis of respiratory behaviors

Respiratory activity was measured using whole-body plethysmography in a room with controlled ambient temperature (22-24 °C) and humidity as described before [22]–[24]. Briefly, awake male mice were placed in a plexiglass chamber (∼ 0.5 L), which was flushed with room air (i.e., 21% O_2_ and <0.03% CO_2_), at a rate of 1.2 L min^-1^ during measurements of baseline respiratory behavior. Concentrations of O_2_ and CO_2_ in the chamber were monitored using a fast-response O_2_/CO_2_ analyzer (ML206, AD Instruments). To account for changes in base physiology due to circadian changes, all experiments were performed between 10:00 and 14:00 hours. The baseline breathing was recorded 40 min after placing the animal in the recording chamber, after which the animal was exposed to hypoxic conditions (10% O_2_) for 5 min. In separate experiments, the same mice were exposed to hyperoxic-hypercapnic conditions (6% CO_2_ and 60% O_2_; 5 min) after a minimum of two weeks following the hypoxic experiment. The hyperoxic condition (i.e., 60% O_2_) in the setting of hypercapnia decreases the effect of the peripheral chemosensitive regions on breathing [25].

### Measurement and analysis of locomotion activities

The mice’s basic locomotion was tested in an open-field arena as described elsewhere [26]. Briefly, mice were placed in a square enclosure (50 × 50cm) to roam freely for 1.5 hours. The cumulative distance traveled and locomotion speed in the open-field apparatus were tracked, recorded, and quantified using ANY-maze behavioral tracking software.

### Measurement and analysis of spontaneous and induced grooming activities

The procedure of self-grooming behavior measurement was adapted from a previously published study [27]. Mice were placed individually into the plethysmography chamber or a clean home cage and allowed to habituate for 20-30 min. *Spontaneous* self-grooming behavior was recorded for 10 min. A timer was used to assess the cumulative time spent in self-grooming behavior, which included paw licking, unilateral and bilateral strokes around the nose, mouth, and face, paw movement over the head and behind the ears, body fur licking, body scratching with hind paws, tail licking, and genital cleaning. The number of self-grooming bouts was also counted. Separate grooming bouts were defined as a pause greater than 5 seconds or the occurrence of behaviors other than self-grooming. Self-grooming microstructure was not assessed. *Induced grooming*: A standard spray bottle was filled with distilled water, and the nozzle was adjusted to the “misting” mode. Mice were placed on a flat surface and sprayed three times from 30 cm away to be adequately covered with mist. Mice were placed individually into the testing enclosure, and grooming behavior was recorded for 10 min following the spray and then analyzed as described in the preceding section.

## RESULTS

### *Gnptab*-mutant mice exhibit impaired respiratory response

We and others have shown that mild metabolic challenges (hypoxia or hypercapnia) affect breathing behaviors in rodents [24], [25], [28]–[38]. Therefore, to characterize breathing behaviors, we exposed *Gnptab*-mutant and control littermates to hypoxia or hyperoxic-hypercapnia and compared respiratory parameters between *Gnptab*-mutant and control littermate mice across three gas conditions: room air (21% O_2_, 0% CO_2_), hypoxia (10% O_2_), and hyperoxic-hypercapnia (60% O_2_, 6% CO_2_). The breathing rate was not different in room air between *Gnptab*-mutant mice and controls (194 ± 7 min^−1^ vs 181 ± 6 min^−1^ in control; *P* = 0.2; *t* test), nor in hypercapnic conditions. However, hypoxia-induced augmentation of breathing frequency was lower in mutants (238 ± 6 min^−1^) than controls (264 ± 10 min^−1^, *P* = 0.032; *t* test; Figure 1A). This blunted response under hypoxic but not hypercapnic conditions indicates that respiratory circuits might be functioning at a lower excitation state in *Gnptab*-mutant animals [24]. On the other hand, tidal volume (i.e., depth of breathing) did not differ between genotypes under any breathed gas conditions (Figure 1B). As expected, minute ventilation, a composite measure of breathing rate and tidal volume, was lower in *Gnptab*-mutant under hypoxia (47 ± 2, control: 57 ± 3, *P* = 0.024; Figure 1C). We also measured the regularity of breathing by calculating the irregularity index [24], [30]. Our data suggest that breathing was less regular (i.e., higher irregularity index) in *Gnptab*-mutant mice than in control mice at room air (48 ± 4 % vs 23 ± 3 min^−1^ in control; *P* < 0.001; *t* test) and under hypoxic conditions (29 ± 2 % vs 19 ± 1 min^−1^ in control; *P* < 0.001; *t* test). Irregular breathing in room air and under hypoxic conditions but not hypercapnic conditions further suggests that the respiratory circuits might function at a lower excitation state [24]. When combined, these results demonstrate that *Gnptab-*mutant mice exhibit irregular breathing, with blunted O_2_ chemoreflex control of breathing.

**Figure 1.**
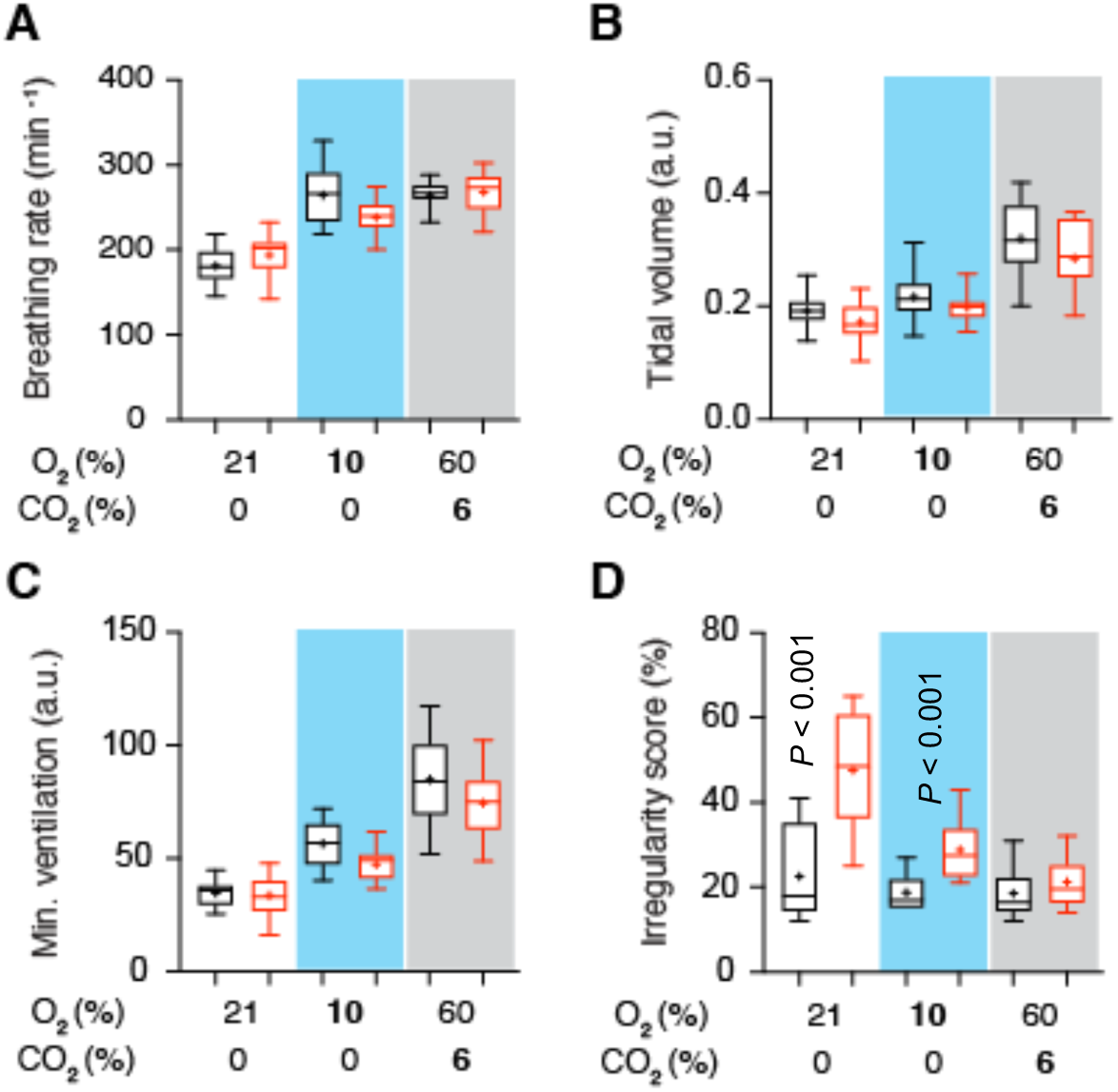
Altered breathing behaviors in *Gnptab*-mutant mice. Respiratory parameters were measured in control (black) and *Gnptab*-mutant (red) adult male mice under normoxic (21% O_2_), hypoxic (10% O_2_; blue shaded area), and hypercapnic (6% CO_2_; gray shaded area) conditions. (**A**) Breathing rate (breaths per minute) increased in both groups under hypoxia and hypercapnia, but mutants showed attenuated responses during hypoxia. (**B**) Tidal volume showed modest increases with gas challenges, especially during hypercapnic conditions, with no major genotype differences. (**C**) Minute ventilation increased in both control and *Gnptab*-mutant mice. (**D**) Irregularity score (%), a measure of breath-to-breath variability, was elevated in *Gnptab*-mutant mice under normoxic and hypoxic conditions, but not hypercapnic conditions. The blue and gray shaded areas represent hypoxic and hypercapnic exposures, respectively. Data sets without *p* values indicated are not significantly different. *P* values – unpaired *t*-test; a.u. – arbitrary units.

### *Gnptab*-mutant mice display delayed and disorganized grooming behavior

Next, we evaluated spontaneous self-grooming behavior in *Gnptab*-mutant and control littermate mice. Latency to the first spontaneous grooming bout was significantly increased in *Gnptab*-mutant mice (9.3 ± 2.5 s) compared to controls (2.3 ± 0.2 s, *P* = 0.02, *t* test; Figure 2A), suggesting impaired initiation of stereotyped motor sequences.

**Figure 2.**
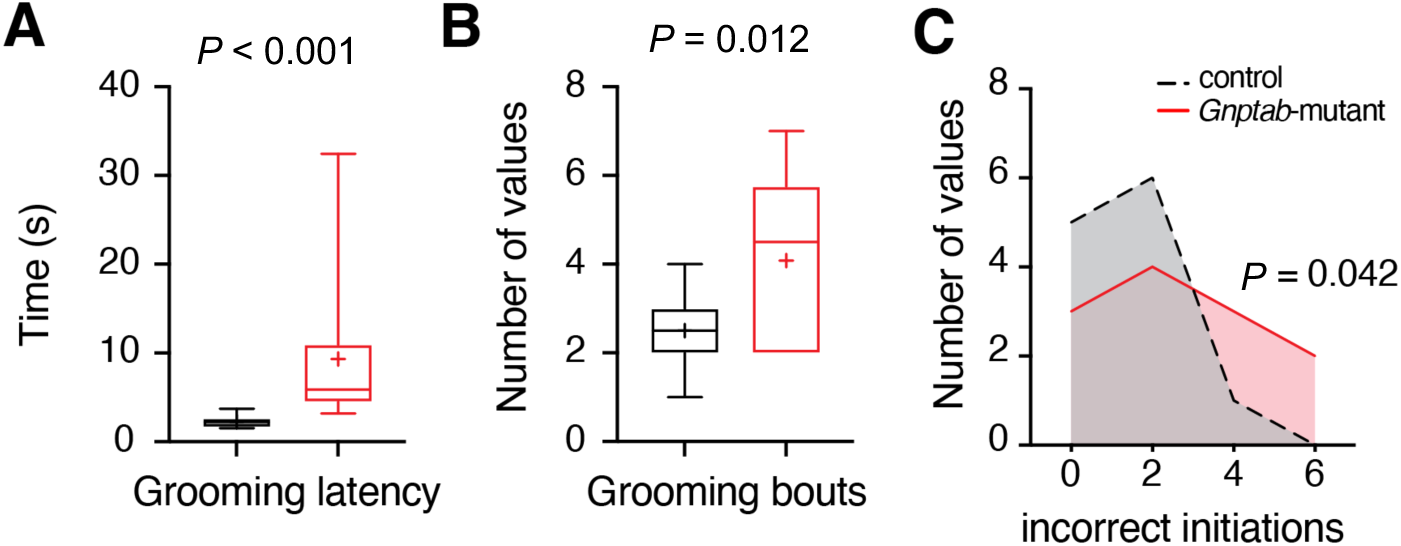
Changes in self-grooming behaviors in *Gnptab*-mutant mice. **(A)** Grooming latency, defined as the time to initiate the first grooming bout, was significantly increased in *Gnptab*-mutant mice, indicating delayed initiation of grooming behavior. **(B)** Grooming bouts, or the number of discrete grooming episodes, were elevated in mutants compared to controls, suggesting increased grooming activity. **(C)** Incorrect grooming initiations, defined as disruptions in the typical grooming sequence, were more frequent in *Gnptab*-mutant mice. *P* values – unpaired *t*-test

Number of grooming bouts was higher in mutants (4.1 ± 2) than in controls (2.5 ± 1, *P* = 0.011, *t* test; Figure 2B), indicating either increased grooming drive or more fragmented sequences. In terms of grooming structure, *Gnptab*-mutants committed more incorrect grooming initiations (defined as out-of-order or incomplete sequences) than control mice (median error count = 2 vs. 1, p = 0.042; Fig. 2C), suggesting an atypical but rigid behavioral organization. Accordingly, it is seen that *Gnptab*-mutant mice have delays or alterations in spontaneous rhythmic behavior.

## DISCUSSION

Stuttering is a neurodevelopmental motor control disorder that affects the pattern and timing of speech initiation, but existing data suggest that its effects extend beyond the vocal domain. Motor differences, including differences in finger sequencing, bimanual coordination, and orofacial nonspeech movements, have been documented in people who stutter [39]–[43][44]–[47][48], supporting the view that stuttering reflects a broader impairment in the initiation or coordination of rhythmic motor behaviors, including but not limited to vocalization. Although the involvement of cortico-basal ganglia-thalamo-cortico loop is proposed, the fundamental circuit and cellular processes are still not well understood.

In this study, we used a genetically defined mouse model [17], [18] to investigate how mutations in *Gnptab*, a gene previously linked to persistent stuttering in humans [49]– [53], impact non-vocal motor systems. Prior work has shown that *Gnptab*-mutant mice exhibit vocalization abnormalities during development [17], [18] and adulthood [20]. However, whether other rhythmic behaviors are disrupted has not been systematically examined.

We found that *Gnptab*-mutant mice exhibit selective impairments in breathing under hypoxic conditions, along with increased breath irregularity in normoxia and hypoxia. These results are consistent with previous work showing that breathing control circuits, particularly under conditions of reduced metabolic drive (i.e., hypoxia), are susceptible to disruptions in excitation–inhibition balance and neuromodulatory signaling [24], [25], [28]. Interestingly, the preserved hypercapnic response in mutants suggests that core central CO_2_ chemoreflex pathways remain functional, but may be operating at a lower excitation threshold, limiting the ability of breathing centers to adapt to hypoxia. This dissociation points to a subtle but functionally meaningful impairment in state-dependent respiratory control, one that may model similar irregularities reported in people who stutter [7]–[11].

The finding that *Gnptab*-mutant mice also show delayed grooming initiation and increased grooming stereotypy extends this low-excitation hypothesis to cortical and basal ganglia circuits involved in internally guided behavior. Grooming is a highly conserved, patterned motor sequence that requires precise timing and behavioral adaptability. The increased number of grooming bouts and elevated rate of incorrect initiations in mutant animals compared to littermates resemble the motor rigidity and sequencing errors observed in human studies of stuttering and related developmental disorders [12] [54]. These behavioral disruptions emerge spontaneously, without external provocation, reinforcing the hypothesis that the *Gnptab* mutation impairs intrinsic motor patterning mechanisms, possibly through astrocyte-specific dysfunction within motor circuits [17], [24], [38], [55]–[57]. Recent evidence indicates that astrocytes in *Gnptab*-mutant mice show regionally specific anatomical impairments [16], and that astrocytic modulation is essential for the proper timing and adaptability of central pattern generators that control rhythmic motor outputs [38].

Together, these findings support a model in which Gnptab-dependent lysosomal trafficking is required for maintaining appropriate excitability and coordination within motor control circuits. While Gnptab is broadly expressed and essential for overall cellular function, its disruption selectively impacts rhythmic motor systems that require precise timing and dynamic regulation. These include vocal motor circuits, respiratory centers, and forebrain pattern generators involved in grooming. The convergence of deficits across these domains strengthens the case for using *Gnptab*-mutant mice as a model of multisystem motor timing disorders and provides a mechanistic framework for understanding the non-vocal symptoms that are linked to this mutation. Further studies should clarify whether these phenotypes result from neuronal or glial dysfunction and whether altered lysosomal signaling affects specific neuromodulatory systems that regulate motor pattern flexibility. This model could help bridge molecular genetic findings and cellular lysosomal pathways, with system-level dysfunctions in the modulation of circuit excitability reported in stuttering and other developmental motor disorders.

## Acknowledgements

We thank the NINDS Light Microscopy Core and the NIMH Section on Instrumentation for technical assistance and resources. We are grateful to Drs. Jeffrey Smith (NINDS), Yogita Chudasama (NIMH), and David Leopold (NIMH) for support and mentorship. We also thank Dr. Dennis Drayna (NIDCD) for providing the mouse model. This work was supported by the Intramural Research Program (IRP) of the NIH, NINDS (ZIA NS009420 to SSB), and in part, by the NIMH IRP Rodent Behavioral Core (MH002952).

